# Viral Complexity: Amino acid co-evolution in viral genomes as a possible metric

**DOI:** 10.1101/159541

**Authors:** C. K. Sruthi, Meher K. Prakash

## Abstract

Viruses are simultaneously simple and complex. Simple because they have barely around ten types of proteins compared to tens of thousands of proteins in bacteria. Complex because amino acid mutation rates are very high, challenging host immune system and drugs. In this work we use the co-evolution of amino acids and the network characteristics that arise out of it to describe the complexity hidden in the multitude of variations in a viral genome. Using large-scale genomic data, the complexity in several viruses was compared. Interestingly, the co-evolutionary relations were primarily intra-protein in avian influenza and inter-protein in HIV-1. The network degree distributions showed two universality classes: a power-law with exponent −1 in HIV-1 and avian-influenza, random co-evolutionary behavior in human flu and dengue, suggesting the co-evolution as one way to statistically classify the complexity in viruses. The observed correlation between the network densities and the strengths on virus Richter scale raises interesting questions on whether it is possible to define the complexity of viruses using their evolutionary networks.

## Introduction

The genome size and complexities in different organisms vary widely. While bacteria have genes encoding several thousand types of proteins, most viruses have barely around ten types of proteins. This is true for viruses as benign as common flu to the lethal ones like ebola. Interestingly, as the number of base-pairs encoding these genes varies from hundreds of millions to tens of thousands, the mutation rate which is the chance of making an error over a generation increases by many orders of magnitude.^1, 2^ Despite this high rate of mutations or errors in the amino acids of viral proteins, many viruses remain functional and infect the hosts possibly because many deleterious mutations are compensated by other simultaneous mutations. Continuously evolving viruses thus become much more unpredictable both for the immune system as well as the drugs developed against them. Characterizing the evolutionary behavior of viruses will thus be an important step towards understanding the complexity of viruses. Yet, to date there is no informatics way of describing the complexity of viruses and their evolution.

One way of describing the biological systems-level complexity involved in healthy and diseased cells is by studying interaction networks. Biological networks can be formed out of transient molecular interactions such as in proteins interacting with other proteins or from persistent physical interactions such as in neural networks. Metabolic^3^ and gene regulatory networks,^4^ protein-protein interaction networks,^5, 6^ and neural networks are examples of functional cellular networks. Disease networks on the other hand try to connect genotypes with phenotypes.^7, 8^ Protein-protein interaction networks have been used to describe the complexity of the different systems from *E. Coli* to humans^9^. Protein interactions became fine grained as the *C. elegans* interactomes initially identified and mapped at protein level^10^, subsequently focused at the domain level.^11^ Since viruses have only around ten types of proteins, but high mutation rates, further fine-graining with a focus on amino acid interactions is statistically more meaningful.^12–15^ Studying the co-evolutionary relations among the amino acids is an important step towards describing and eventually deciphering the complexity of the viruses.

Amino acid level co-evolutionary interactions can arise either from structural constraints between proximal amino acids or because of functional constraints from amino acids at distal sites or other proteins. Several studies focused on amino acid interaction networks, starting from the three dimensional structural data of the proteins.^12, 13, 16^ The utility of structure based methods is limited because of the limited structural information available, as well as because it more likely highlights the proximal relations. Conversely, using amino acid co-evolutionary couplings from abundant homologous sequence data of multiple species,^17^ bioinformatic approaches such as Statistical Coupling Analysis (SCA)^18^ and Direct Coupling Analysis^19^ could predict hotspots of proteins, active centers of enzymes, to make *de novo* three dimensional structure prediction of proteins^20, 21^, to identify functionally related clusters of amino acids^22^ and predict the vulnerability of viruses.^23^ In studying viruses where there could be coupled relations between multiple proteins, it is thus useful to explore this functional coupling. In this work, we use large-scale complete genome data to build and analyze amino acid co-evolutionary networks. The data is further analyzed to identify patterns or randomness in this co-evolution in intra‐ and inter-protein amino acids. The complexity of the different viral co-evolutions is also compared by studying the robustness of the network to a targeted or random removal of nodes from the network.

## Results and Discussion

### Data inclusion

Complete genome sequence data were obtained from the NCBI servers. The analysis was performed when large scale genomic data, from at least 1000 patients, was available. With the current publicly available data, only five viruses were chosen for analysis: HIV-1, hepatitis B, dengue, avian and human influenza. Our analysis was performed on data sets from minimum of 1,784 patient data (HIV-1) to about 8,689 (human influenza). However, the availability of such data is increasing, and in this work we focus on questions that can be posed with such large scale genomic data. Multiple Sequence Alignment (MSA) of the whole genome data from all patients was performed. Using a consensus sequence as a reference, the entire MSA was converted into a binary representation, 1 if the amino acid at a given position in a sequence is the same as that in the consensus sequence, 0 otherwise. Using the Statistical Coupling Analysis protocol,^18^ weighted co-evolutionary matrix **C** that quantifies the relations among the different amino acids was created (**Methods** section). The data on pairwise co-evolutionary couplings was represented using a network for better visualization and analysis. The network representation translates the co-evolutionary information into *nodes* (amino acids) and *network edges* (connections between the amino acids) if the co-evolution matrix element *C_ij_* relating amino acids *i* and *j*, is more than a threshold *C*, *C_ij_ > C^th^*.

### Clustering

Using the complete genome data from different patients, the co-evolution networks for different viruses were constructed. Clustering of nodes was performed using correlation as a weight (**Figure 1**) with the goal of observing patterns which are more general than those seen in pair-wise relations. About 3 to 4 significant clusters can be seen in each of the viruses, and no significant differences in the number of clusters were found when we performed Principal Component Analysis and used Cattell's criterion. However, there is a noticeable difference in the composition of each of the clusters in different viruses. Each of the clusters in the network of HIV-1 co-evolution network have a mixed representation from multiple proteins, suggesting a strong evolutionary relation across the genome, while avian influenza clusters are mostly from intra-protein relations. The inter-protein co-evolutionary relations are much stronger in HIV-1 (**Supplementary Tables 1a, 1b**).

**Figure 1.**
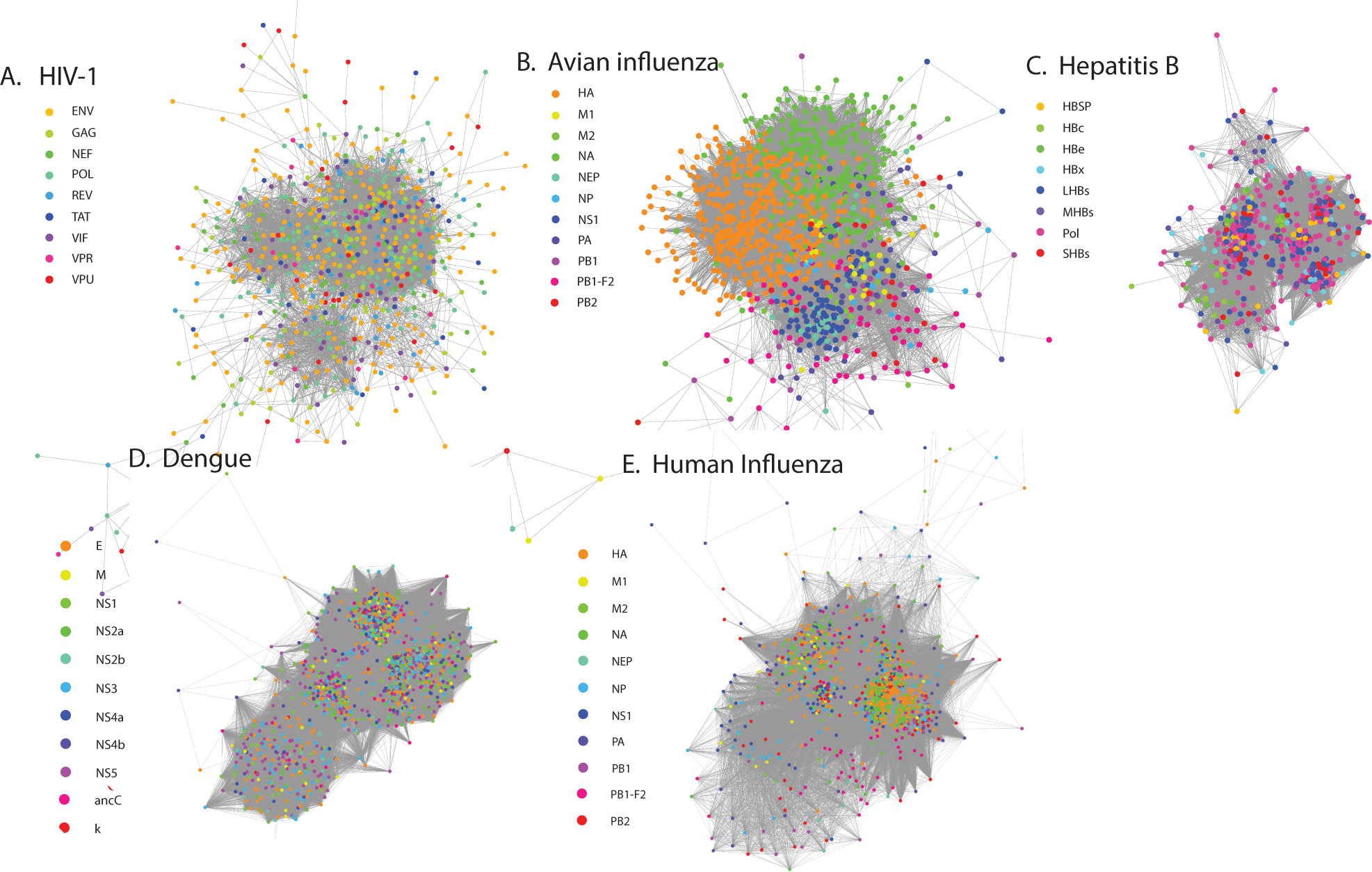
Co-evolutionary network from complete genome analysis of different viruses: (a) HIV-1, (b) avian influenza and (c) Hepatitis B (d) Dengue (e) Human influenza. The networks are generated using co-evolution strength as a weight. The side bar indicates the different types of proteins found in these viruses, as well as the coloring notation used. The networks show three to four major clusters. While in HIV-1, each cluster has a mixed representation from all the proteins, avian influenza clusters are mainly from intraprotein co-evolutionary relations. Network representations were generated using Cytoscape^31^

### Scale-free vs. random networks

The complexity of the networks is analyzed by studying its node-degree distribution, *n*(*k*) - the number of times a node with a certain number of edges *k* appears in the network.^24^ Two commonly seen universality classes in these distributions - power-law and Poissonian, suggesting systematic or random underlying basis, occur in the amino acid degree distributions as well. In HIV-1, as well as in avian influenza, a power-law *n*(*k*) ~ *k*^−1^, while dengue, human influenza show a Poissonian distribution (**Figure 2**).^24^ Consistent with the observation in the clustering, using only the inter-protein co-evolution from HIV-1 did not change the observed powerlaw. Hepatitis B on the other hand showed a mixed behavior including both power law and Poissonian behaviors (**Figures 4a-4f**). We further analyzed the role of the threshold by varying *C^th^* in the analysis of Hepatits B. As shown in **Figure 4**, as the *C^th^* increases from 1.0 to 3.0, the power-law component becomes more pronounced (similar data for other viruses is shown in **Supplementary Figures 1** to **4**). The data shows a clear separation of network connections arising from two different origins, an organized network of co-evolution above a certain threshold and random network connections at lower thresholds of co-evolution. Within this power-law regime a further change in cutoff did not result in a change in the exponent significantly. The analysis presented so far is the statistical description of data collected from patients and is averaged over all the years of sample collection. In order to study the temporal evolution patterns, we performed time analysis on the data set which is most abundant, human influenza. We divided the complete genome data from human influenza into periods where the number of data sets is similar (~2000 complete genomes each). A node-distribution analysis shows that over this period, there is no significant change in the co-evolutionary complexity of viruses (**Supplementary Figure 5**).

**Figure 2.**
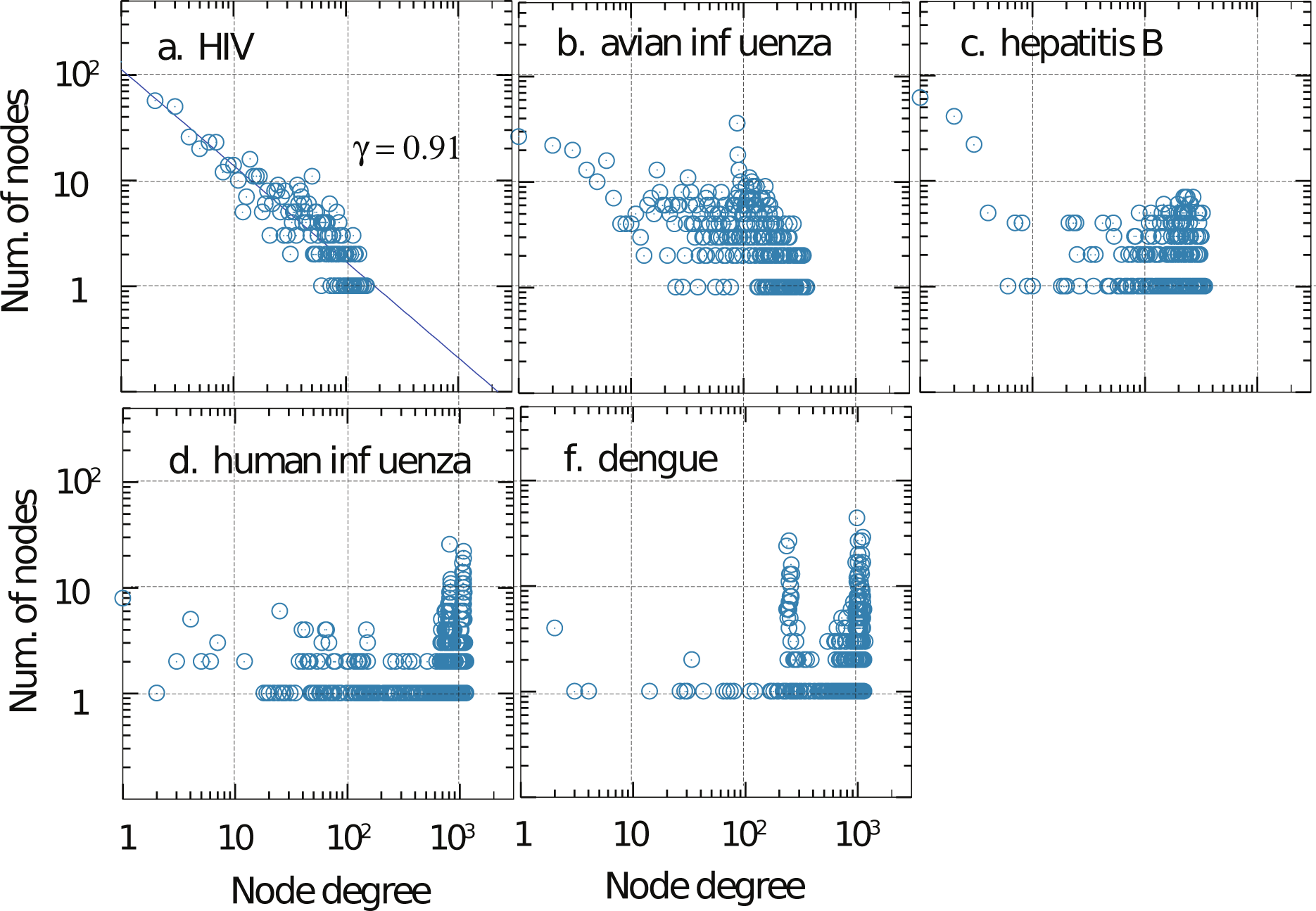
Node degree distribution from the complete genome data in different viruses showing a range of behavior from a pure power-law (HIV-1 and avian influenza) to a pure-random network behavior (dengue and human influenza). A cutoff *C^th^* = 0.85 was used as a threshold for establishing network edge connections. The effect of changing the cutoff is discussed separately.

### Complexity Measure

It is difficult to describe complexity, and even more to quantify it with one single measure. The lack of a simple and precise metric for complexity is a problem both in biology and in network science. For biological complexity of viruses, here we use the strength on virus Richter scale^25^ as a measure of their complexity. While it is understood that the Richter scale indicates mortality from viruses, which includes several factors from how fast the virus mutates to how poor the public health provisions are, for lack of a better way to compare the strengths of viruses or difficulty of developing vaccines against them, we use Richter scale. Figure 3 shows a plot between the virus strength and the network characteristic - network density. Avian influenza data from avian host was not part of this analysis as the Richter scale definition is irrelevant. Interestingly Figure 3 shows a correlation between the network metric and the biological metric. Clearly this correlation is not conclusive, as they are based on studies of just four viruses. However, it raises the possibility that the complexity of the biology and the pathogenicity of the virus may be reflected in the amino acid co-evolution networks.

**Figure 3.**
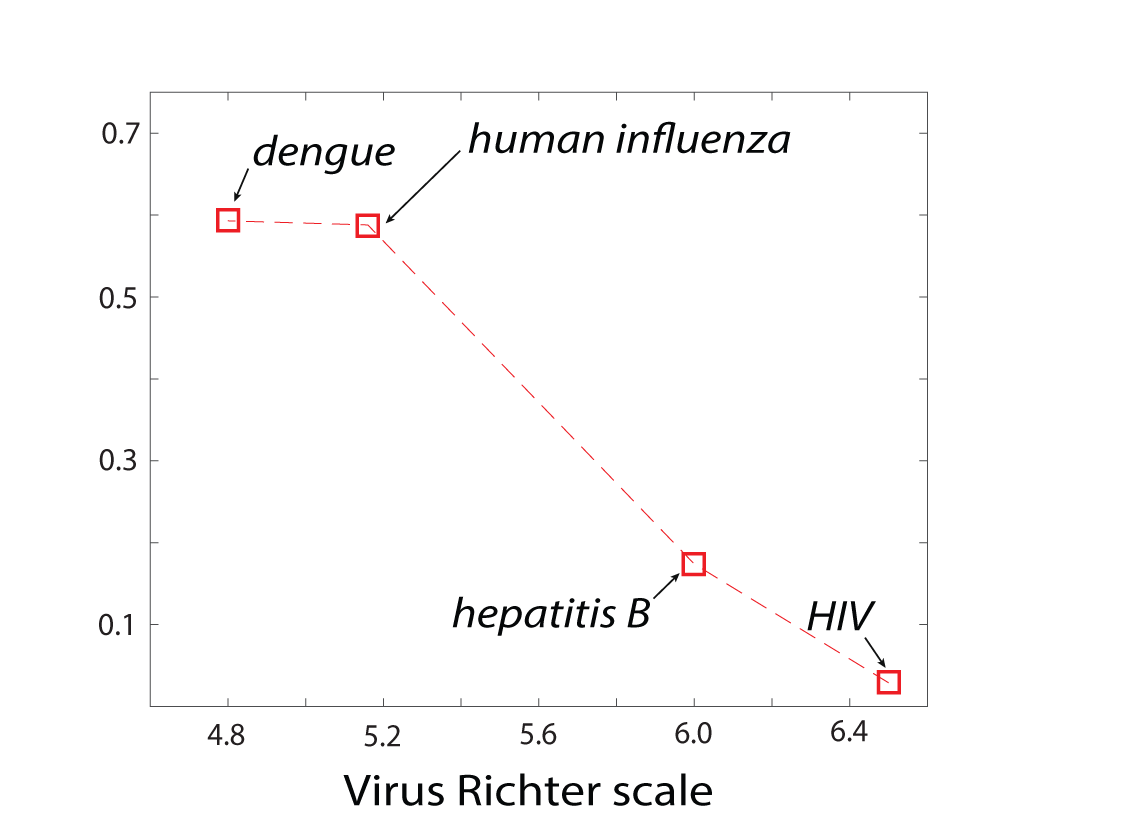
The relation between the complexity of the virus, as described by Virus Richter scale, and network characteristics - node density (□)

**Figure 4.**
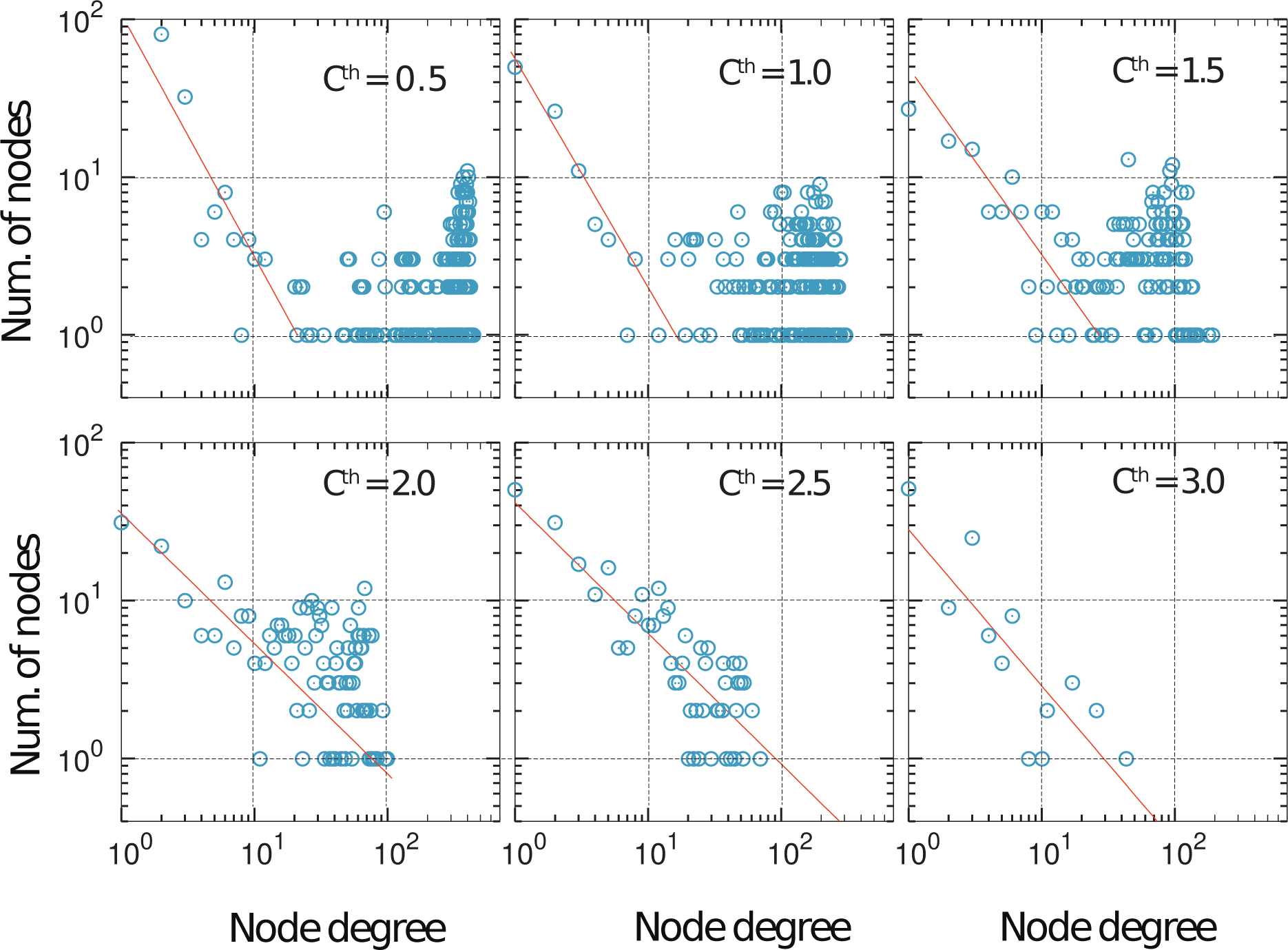
Node degree distribution sensitivity was studied in Hepatitis-B network by changing the cut-off value used for defining edge connectivity between the nodes. At a very low cut-off there is a mixed behaviour in the node degree distribution, with both power-law as well as a random component. As the cutoff is increased, the random component is selectively removed, while preserving the power-law component. This suggests a clear separation of network connections from random and systematic origins. By choosing a threshold value, one can filter and study just the systematic component.

### Power law

Random networks (Erdos-Renyi model), small world networks (Watts-Strogatz model^26^) and self-similar networks (Barabasi-Albert model^27, 28^) arising in diverse contexts such as WWW, protein-protein interactions, co-authorship networks, etc have been well studied. Some of the mechanisms that explain the observed phenomena are preferential attachment model where newer edges are added to a node depending on its current degree, or based on its pre-defined fitness or a potential for a degree. The power-law with *γ* ∼ 1 observed in the co-evolution network is different from the typical power-laws *γ* varying from 2 to 3 and is closer to the behavior in co-authorship networks. Unlike a citation network, there is no reason to believe that the co-evolutionary network evolves with a continuous increase in the number of nodes and edges. Considering amino acid conservation (*ϕ*) as a surrogate for their fitness, we developed a fitness based model^29^. The model uses two distributions derived from the whole genome data: (a) the distribution of the conservation among the amino acids, *p*(*ϕ*) (**Supplementary Figure 6**) (b) the co-evolutionary fitness potential of the node *η*(*ϕ*) corresponding to a given conservation of the amino acids. The latter can be modeled as a gaussian distribution, with minimal co-evolutionary fitness for amino acids with very high and very low conservation, a peak in between at *ϕ_m_* and standard deviation *σ*. Considering a pair of amino acid nodes *i* and *j*, and two random numbers *r*_1_ and *r*_2_ drawn from a uniform distribution, edge *i* — *j* is created if *r*_1_ *r*_2_ *ϕ*(*ϕ_i_*) *ϕ*(*ϕ _j_*). This algorithm generates a node-degree distribution with *γ* ∼ 1 (**Supplementary Figure 7**). For example, for HIV-1, the conclusion is relatively invariant for a gaussian with *ϕ_m_ =* 0.6 0.7 and *σ =* 0.02−0.07. As the parameters go out of this range, node degree distribution eventually transforms to a random network model. While the model captures the observed power-law and poissonian distributions with minimal assumptions, the assumptions need to be related to the evolutionary stages of the viruses to see if the statistical complexity of viral co-evolution can be related to their biological complexity.

### Robustness of networks

The co-evolution networks were checked for their robustness by removing different fractions of nodes and all the edges connecting to them,^30^ the spirit being that the critical amino acids or groups of them can be a potential drug target. The nodes to be removed were chosen either randomly or by picking those with the highest degree, to simulate a random error or a targeted attack, **Figure 5** shows how the effective diameter - a metric of network connectivity - is affected by the targeted or random removal. Interestingly, random removal has the highest impact on human influenza network, and the least affected is HIV-1. Further, the impact of targeted removal is highest on HIV-1. The overall characteristics of robustness can be intuitively expected from the the power-law distribution of nodes. We also used another measure of robustness, which is the number of clusters it breaks into. The conclusions from these calculations, shown in **Figure 5b** are the same as from node removal.

**Figure 5.**
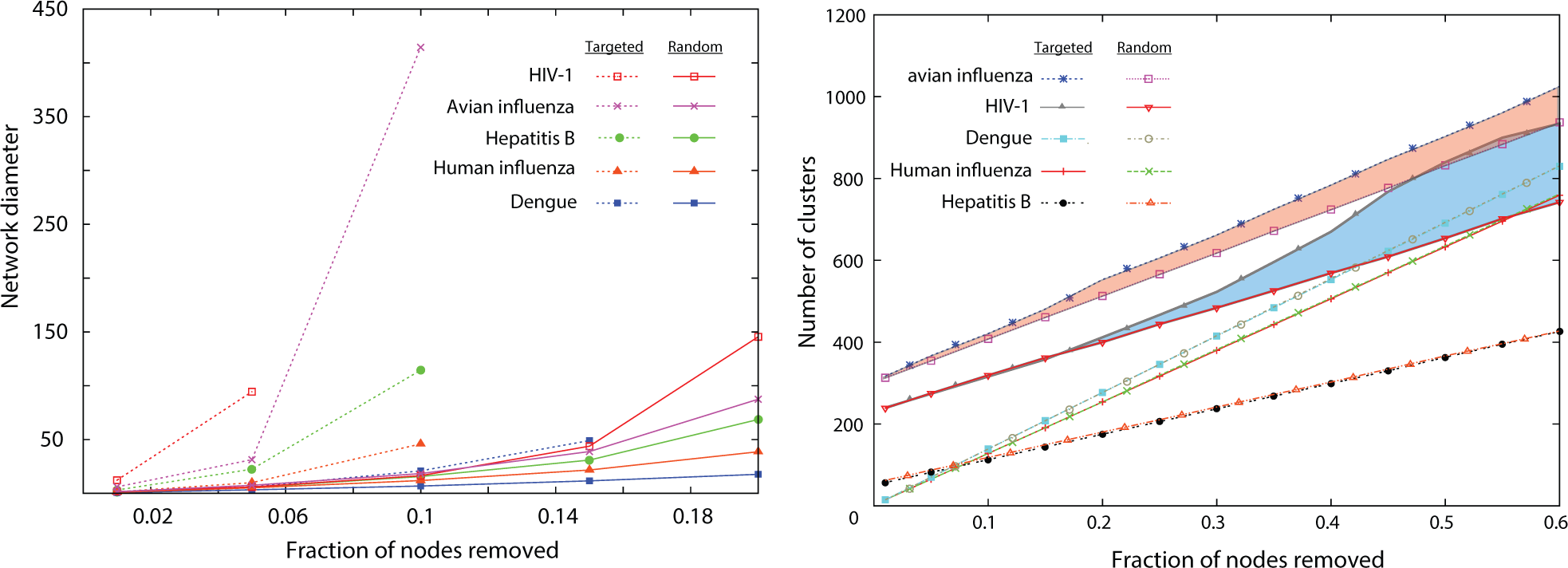
The robustness of the networks is studied using a targeted and random removal of a fraction of nodes. Two different measures were used to estimate how robust the networks are to random and targeted removal: (a) diameter of the network after the removal of the node (b) number of clusters the network breaks into. In HIV-1 and avian influenza there is a clear difference between targeted and random removal, while in others there is not. The avian-influenza and HIV-1 data were shifted up along y-axis by 300 and 200 units for clarity of representation. Network diameter was calculated following the procedure in Ref.^32^

## Conclusions

By using a network representation of amino acid co-evolution we have seen two different characteristics in the large scale complete genome data - clustering with mostly intra-protein or inter-protein couplings and node degrees which have a structured power-law or random origins. When genomic data from more viruses will be available, it will be interesting to see if these two different measures of statistical complexity of genomes can be used to classify viruses into different categories, with a possible mapping to their biological or pathogenic complexity. Further it will be interesting to see if the inter-protein or intra-protein couplings are related to the host adaptation (HIV-1) or the host being a neutral carrier (avian influenza) and how such patterns evolve with time as the viruses adapt from being pandemics to epidemics.

## Methods

### Undirected co-evolution networks

The chance of co-evolution *C_ij_* between a pair of amino acids *i* and *j* is calculated by averaging the columns *i* and *j* of the boolean sequences using either an unweighted or weighted protocol following the Statistical Coupling Analysis protocol.^22^ Unweighted and normalized co-evolution is defined as 
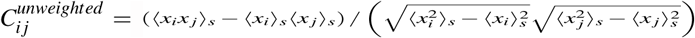
, where *x_i_* is the *i^th^*column in the boolean sequence and 〈〉*_s_* denotes the average over sequences. Weighted co-evolution is defined as 
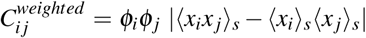
, where 
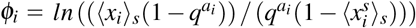
, and *q*^*a_i_*^ is the probability with which the amino acid *a_i_* at position *i* in the consensus sequence occurs among all proteins. background probability of the most frequent amino acid *a_i_* at position *i* frequency of occurs among all proteins. One could work with either 
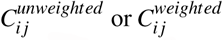
, and in the present work on networks we use 
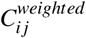
. If the chosen *C_ij_* exceeds a chosen cutoff *c*, we consider an undirected network link *i* — *j* to be present. The sensitivity of the analysis to *c* is discussed in the article. The analysis reported in the article is based on 
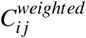
. However, changing the 
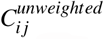
 the power-law distribution in HIV-1 was still around 1, changed from 0.91 to 1.37. Thus, we believe the general conclusions do not change with the weighting.

## Data Availability

The datasets generated and analysed during the current study are available from the corresponding author on request.

## Acknowledgements

We thank Prof. Reka Albert and Prof. Hemalatha Balaram for insightful discussions.

## Author contributions statement

C.K.S.developed Python scripts, performed calculations, data analysis, and literature review; M.K.P. conceived the study, interpreted results and wrote the manuscript.

## Additional information

### Methods

#### Undirected co-evolution networks

The chance of co-evolution *C_ij_* between a pair of amino acids *i* and *j* is calculated by averaging the columns *i* and *j* of the boolean sequences using either an unweighted or weighted protocol following the Statistical Coupling Analysis protocol.^22^ Unweighted and normalized co-evolution is defined as 
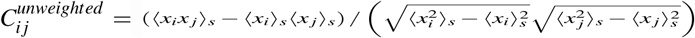
, where *x_i_* is the *i^th^*column in the boolean sequence and 〈〉*_s_* denotes the average over sequences. Weighted co-evolution is defined as 
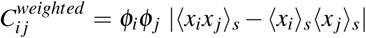
, where 
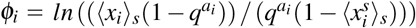
, and *q*^*a_i_*^ is the probability with which the amino acid *a_i_* at position *i* in the consensus sequence occurs among all proteins. background probability of the most frequent amino acid *a_i_* at position *i* frequency of occurs among all proteins. One could work with either 
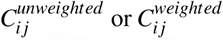
, and in the present work on networks we use 
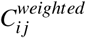
. If the chosen *C_ij_* exceeds a chosen cutoff *c*, we consider an undirected network link *i* — *j* to be present. The sensitivity of the analysis to *c* is discussed in the article. The analysis reported in the article is based on 
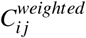
. However, changing the 
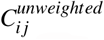
 the power-law distribution in HIV-1 was still around 1, changed from 0.91 to 1.37. Thus, we believe the general conclusions do not change with the weighting.

---

**Supplementary Table 1a.** Table showing the number of inter-protein and intra-protein amino acid co-evolutionary couplings from HIV-1 data, with a *C^th^* = 0.85

| Protein<br>(Num of amino acids) | GAG<br>(500) | POL<br>(1003) | VIF<br>(192) | VPR<br>(96) | TAT<br>(100) | REV<br>(116) | VPU<br>(82) | ENV<br>(856) | NEF<br>(205) |
| --- | --- | --- | --- | --- | --- | --- | --- | --- | --- |
| GAG | 343 | 735 | 287 | 105 | 178 | 248 | 130 | 847 | 277 |
| POL |  | 530 | 398 | 141 | 241 | 319 | 174 | 1112 | 400 |
| VIF |  |  | 93 | 66 | 79 | 126 | 60 | 432 | 143 |
| VPR |  |  |  | 6 | 31 | 43 | 16 | 137 | 50 |
| TAT |  |  |  |  | 25 | 99 | 37 | 297 | 89 |
| REV |  |  |  |  |  | 65 | 55 | 410 | 126 |
| VPU |  |  |  |  |  |  | 16 | 208 | 66 |
| ENV |  |  |  |  |  |  |  | 714 | 462 |
| NEF |  |  |  |  |  |  |  |  | 126 |

**Supplementary Table 1b.** Table showing the number of inter-protein and intra-protein amino acid co-evolutionary couplings from avian influenza data, with a *C^th^* = 0.85

| Protein<br>(No. of amino acids) | NP<br>(498) | PB2<br>(759) | HA<br>(566) | M1<br>(252) | M2<br>(97) | NA<br>(469) | NS1<br>(230) | NEP<br>(121) | PA<br>(716) | PB1-F2<br>(90) | PB1<br>(757) |
| --- | --- | --- | --- | --- | --- | --- | --- | --- | --- | --- | --- |
| NP | 49 | 101 | 219 | 210 | 99 | 77 | 291 | 59 | 137 | 156 | 150 |
| PB2 |  | 56 | 304 | 185 | 78 | 134 | 267 | 49 | 145 | 127 | 130 |
| HA |  |  | 20000 | 900 | 192 | 5362 | 852 | 106 | 184 | 377 | 242 |
| M1 |  |  |  | 171 | 172 | 233 | 570 | 94 | 296 | 268 | 252 |
| M2 |  |  |  |  | 38 | 33 | 233 | 29 | 130 | 103 | 116 |
| NA |  |  |  |  |  | 12005 | 352 | 48 | 30 | 93 | 92 |
| NS1 |  |  |  |  |  |  | 2832 | 1436 | 400 | 352 | 376 |
| NEP |  |  |  |  |  |  |  | 187 | 53 | 66 | 56 |
| PA |  |  |  |  |  |  |  |  | 153 | 168 | 197 |
| PB1-F2 |  |  |  |  |  |  |  |  |  | 637 | 211 |
| PB1 |  |  |  |  |  |  |  |  |  |  | 86 |

**Supplementary Figure 1.**
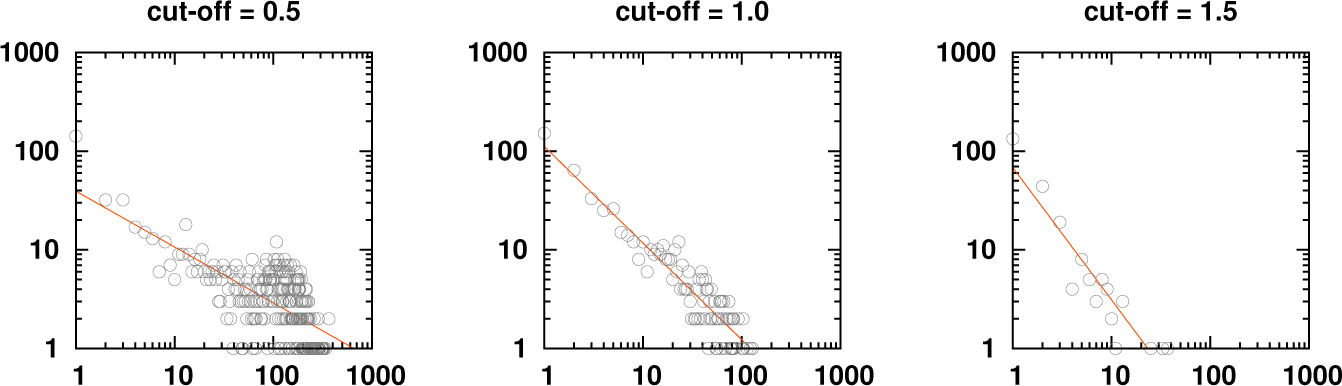
Variation in node distribution of HIV-1 co-evolution network as the cutoff *C^th^* is changed.

**Supplementary Figure 2.**
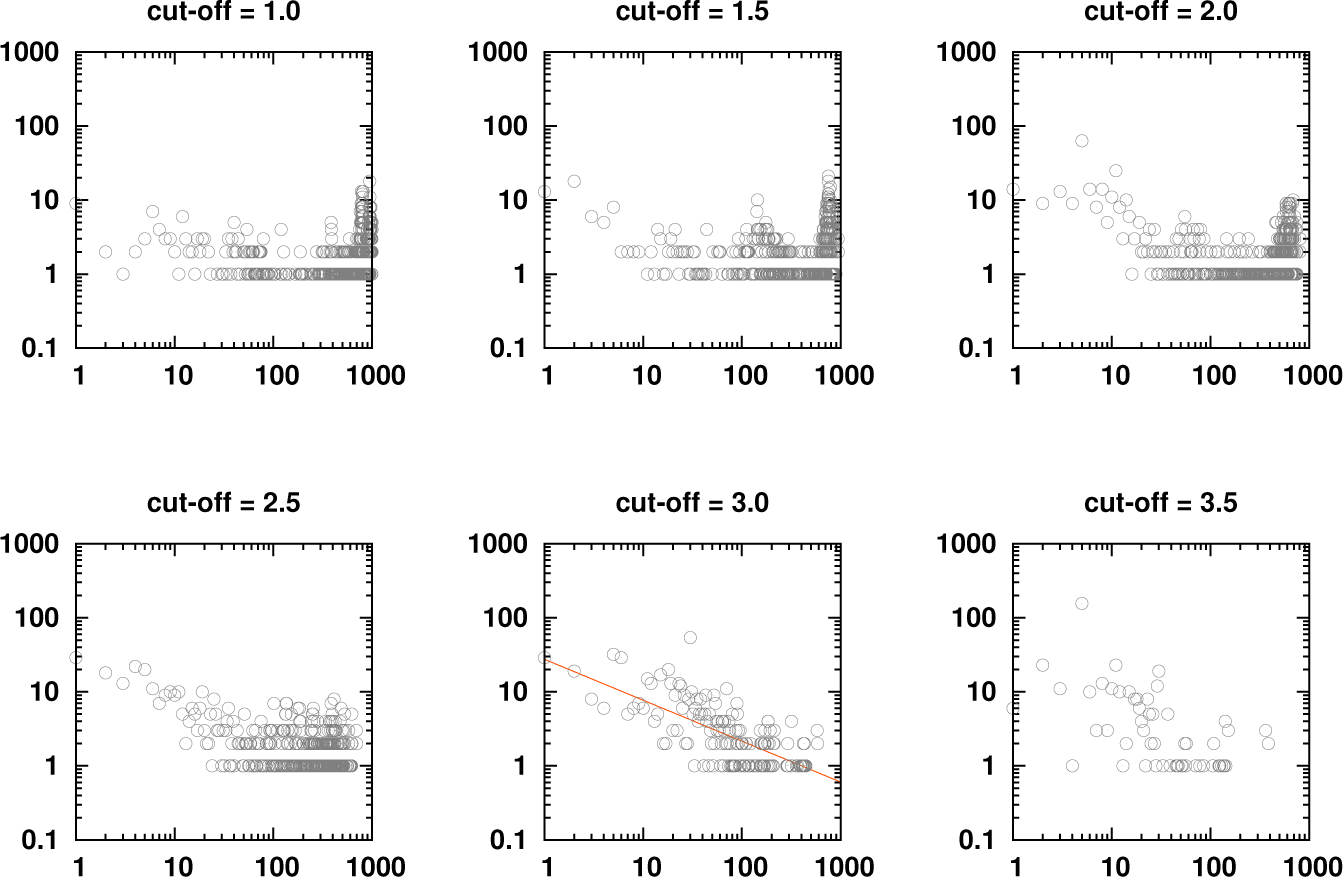
Variation in node distribution of human influenza co-evolution network as the cutoff *C^th^* is changed.

**Supplementary Figure 3.**
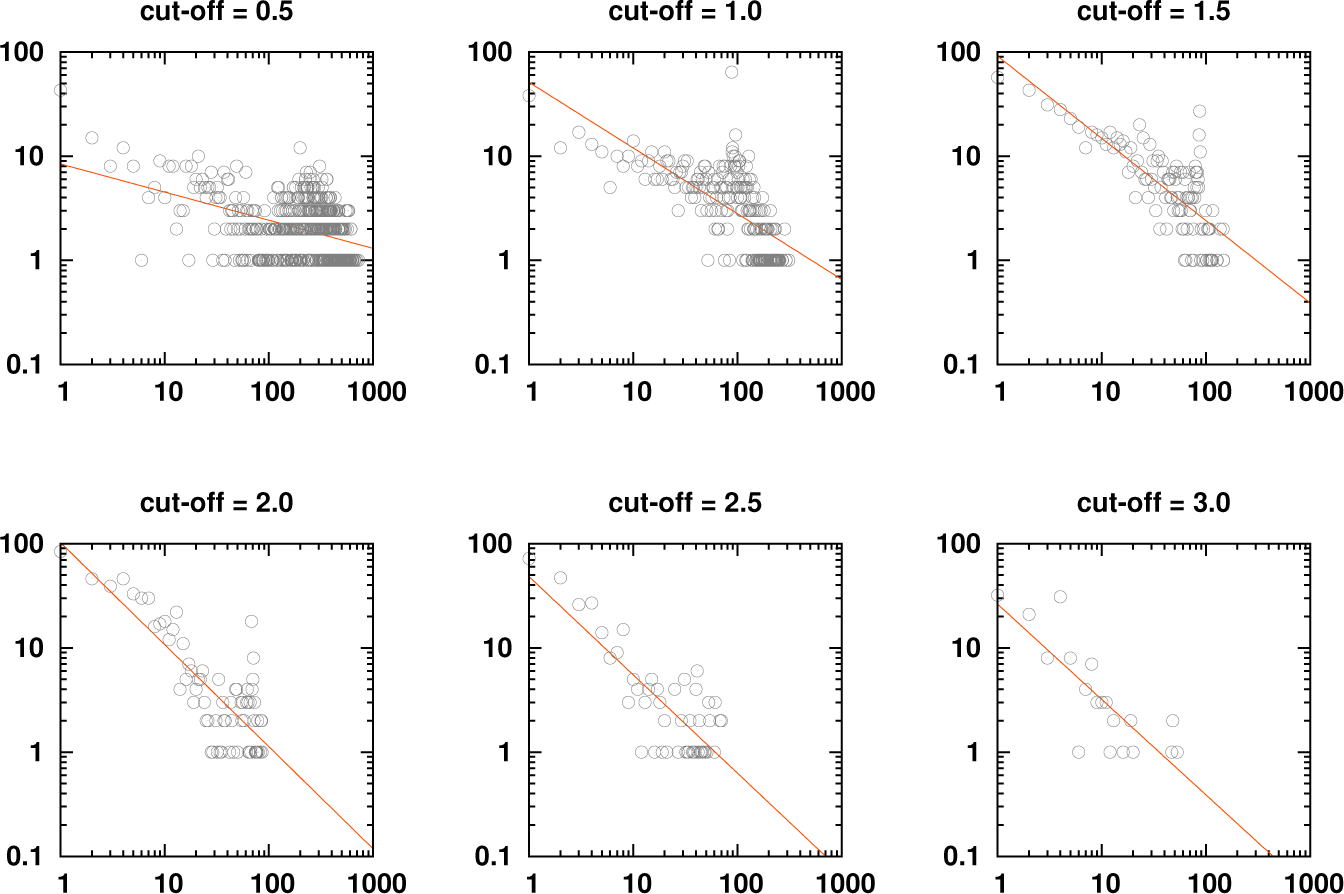
Variation in node distribution of avian influenza co-evolution network as the cutoff *C^th^* is changed.

**Supplementary Figure 4.**
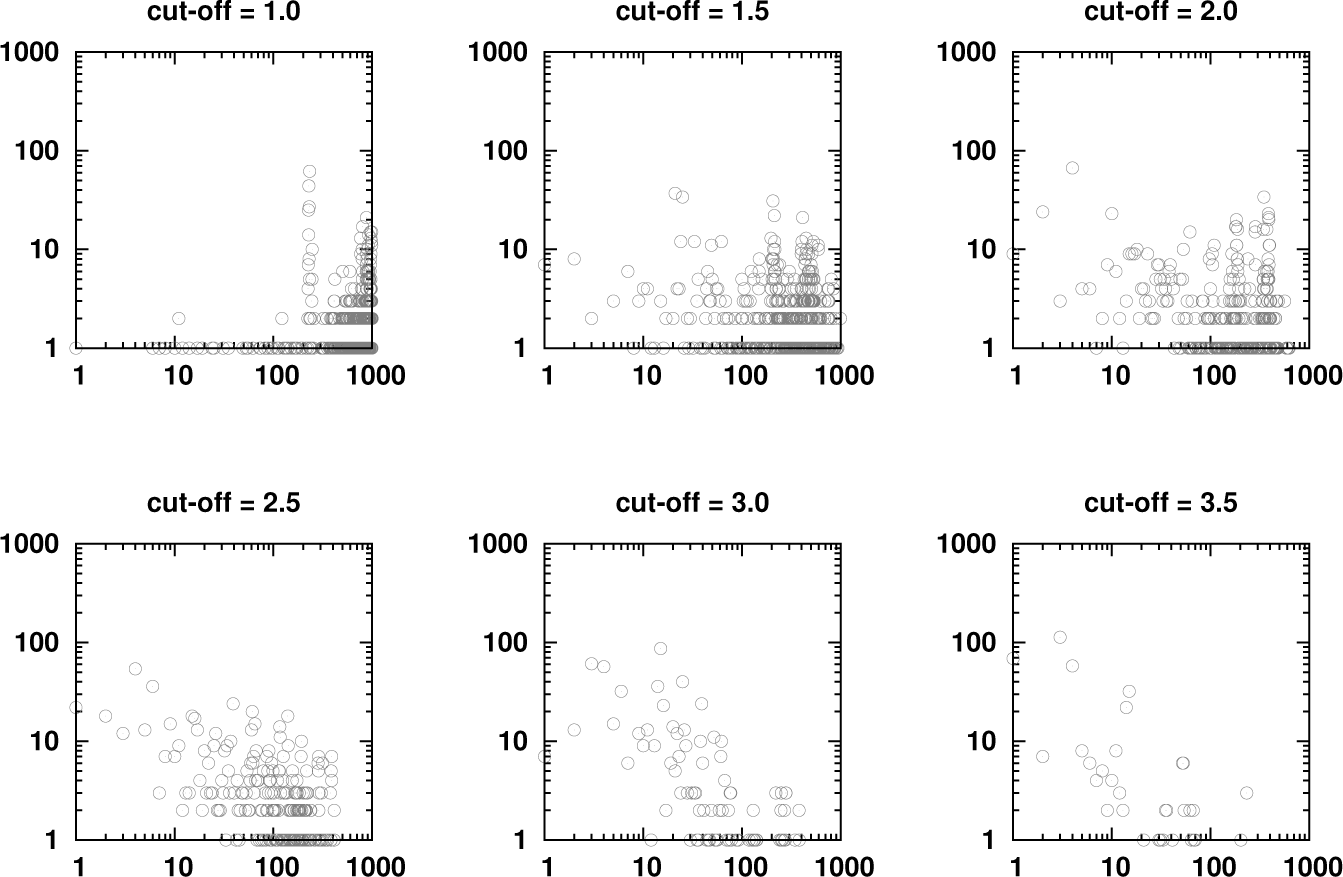
Variation in node distribution of dengue co-evolution network as the cutoff *C^th^* is changed.

**Supplementary Figure 5.**
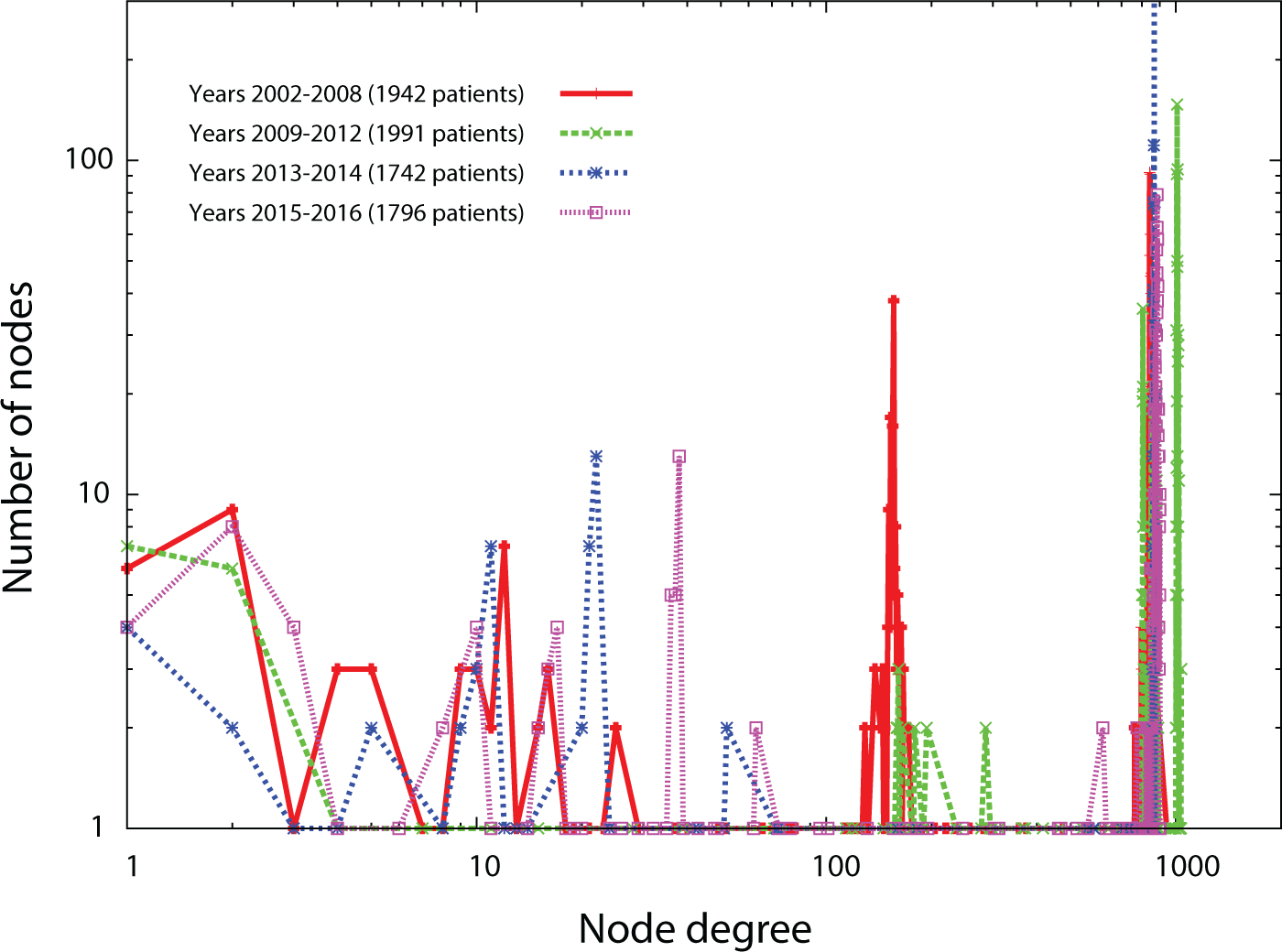
Variation of the node degree distribution over years in the human influenza. Human influenza data was abundant, so we sorted it according to the year of incidence, and made 4 groups of about 2000 patients each. No noticeable trend in the node degree distribution was observed in the data between 2002-2016.

**Supplementary Figure 6.**
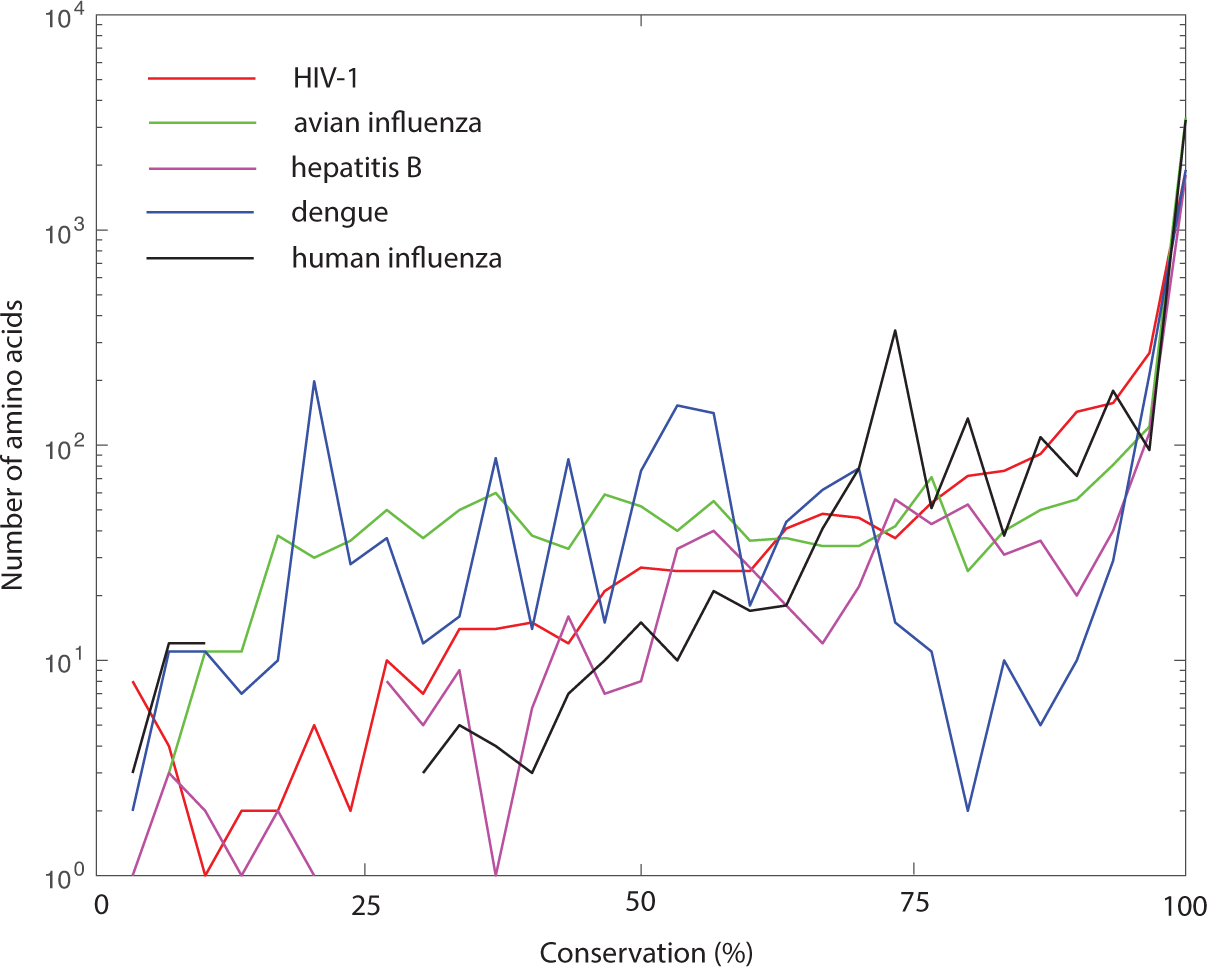
Distribution of the conservation of amino acids in different viruses.

**Supplementary Figure 7.**
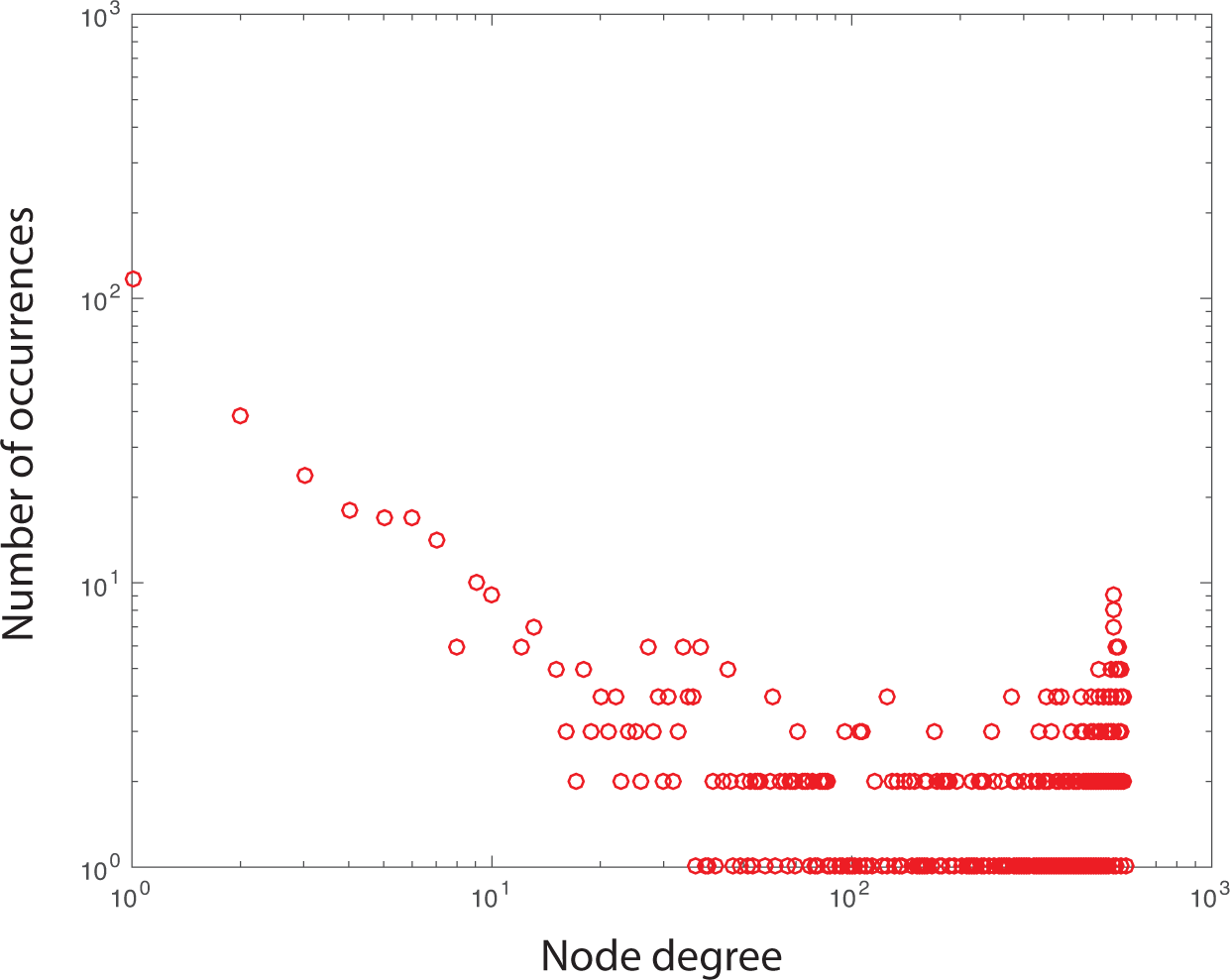
Model network generated using the amino acid conservation distribution from HIV-1, and *η*(*ϕ*) with parameters *ϕ_m_* = 0.05 and *σ* = 0.7

